# Fine Sieving of Collected Atmospheric Particles using Oil Electrophoresis (iSCAPE)

**DOI:** 10.1101/2020.01.04.894998

**Authors:** Xinyue Li, Siyu Xu, Maosheng Yao

## Abstract

It is rather challenging to separate atmospheric particles from nano-to micro-metre mixed in a sample. Here, a system named iSCAPE was invented to efficiently sieve particles out from a mixture by employing an electrostatic field and a non-conductive mineral oil. Tests with atmospheric particles of different cities as well as soil and road dust samples demonstrated that the iSCAPEd particles under different operating conditions moved rapidly with different velocities and both directions. Particles of different sources such as ambient air, soil or road were shown to have different polarity-charged particle fractions, and exhibited clearly different particle electrical mobility graphs after the iSCAPE sieving from seconds to minutes. Data also revealed that after the sieving some particles were enriched at specific mobility ranges. Bacterial ATP measurements implied that the iSCAPE can be also used to efficiently separate bacteria of different sizes and charge polarity. Experimental data here suggest that the iSCAPE sieving strongly replies on the electrostatic field strength, mineral oil viscosity and the run time. In theory, the iSCAPE system can be used to extract any desired targets from a complex sample, thus opening up many outstanding opportunities for environmental, biomedical and life science fields.

**TOC:** 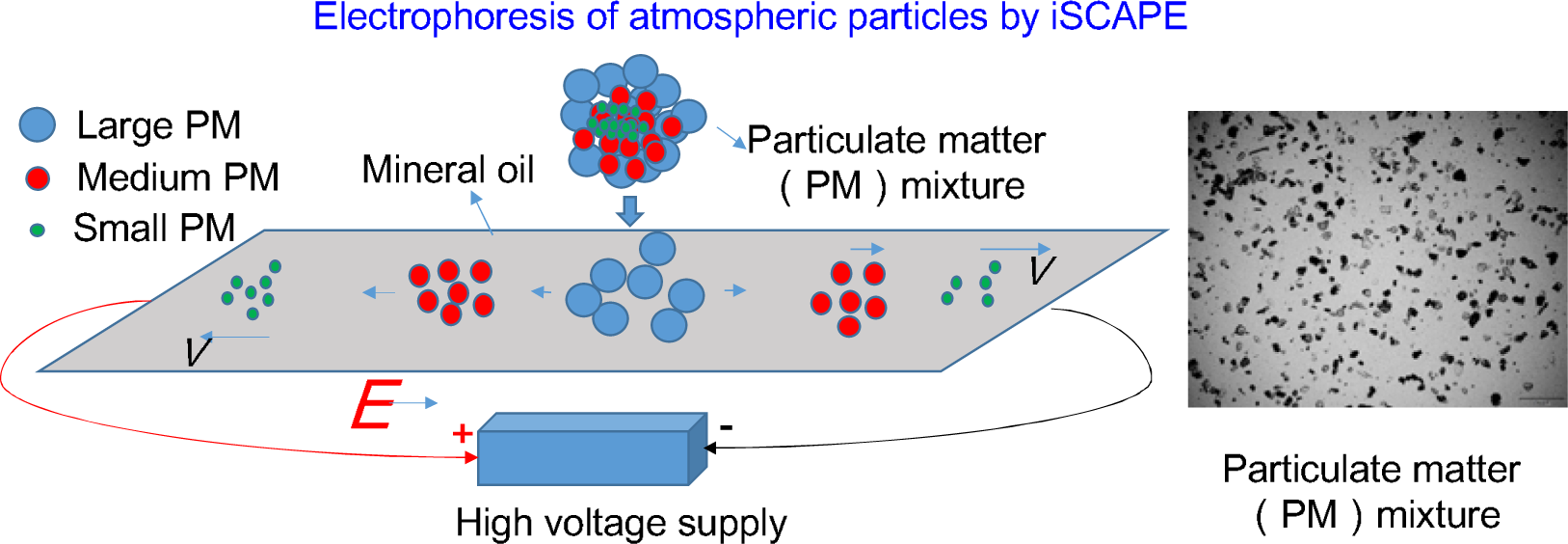

## Introduction

Air pollution, especially the particulate matter (PM), has become one of the most important environmental problems in the world. Exposure to PM has resulted in millions of deaths globally *(1)*. The components of atmospheric particles are very complex, including biological such as bacteria, fungi, viruses, pollen and chemical components - sulfate, nitrate, ammonium and other non-biological particles *(2)*. There are some commercially available instruments for studying size distributions of atmospheric particles *(3–4)*, but they do not automatically provide samples for post analysis. In addition, collection of nanoscale particles requires expensive equipment and high power source *(5)*. Differential mobility analyzer (DMA) with up to 192 size channels is otherwise used to study size distributions of nanoscale particles (1 nm-1μm) *(6–7)*. In addition to its limited size ranges, it is also difficult to collect enough nano-sized particles for post analysis due to its small size and low flow rate. For studying PM health effects, it is also challenging to differentiate the toxicity between different particles since they are often mixed together in a sample. Separation and classification of atmospheric particles using currently available methods are often restricted in terms with their sizes and species, especially for post-analysis.

On the other hand, biological detection of certain microbial species is often prohibitive due to complex environmental matrix of the samples, e.g., PCR inhibition problems encountered in many studies *(8–10)*. In microbiology field, the method of gel electrophoresis has been extensively used in separating the DNAs since its earlier invention *(11–12)*. On another front, it was revealed that particles in the atmosphere likewise carry different polarity charges and levels *(13–15)*. For example, it was shown that bacterial particles in indoor and outdoor air carried about 21-92 elemental unit charges *(15)*. Here, we invented a novel particle mixture sieving system named iSCAPE by employing an electrostatic field together with a non-conductive mineral oil medium. Under the same operating conditions, particles in a sample with different electrical mobility would move at varying velocities in the mineral oil, thus ending up at different locations on the particle moving line within a given time. Depending on objectives, targeted particles or molecules can be thus efficiently extracted from a complex particle of environmental or medical origin using the iSCAPE developed.

## Materials and Methods

### Experimental setup

In this work, we pioneered a novel system named iSCAPE (f**i**ne **S**ieving of **C**ollected **A**tmospheric **P**articles using Oil **E**lectrophoresis) by using an electrical field together with a non-conductive liquid (mineral oil) as shown in Fig. 1. The iSCAPE sieves particles of different sizes from collected atmospheric particle mixture based on their electrical mobility difference. The system consists four major components: high voltage supply (BertanTM, Model 205B-20R, Hicksville, New York), two copper electrodes, mineral oil (M5904, Sigma-Aldrich, USA), and the electrophoresis container (electrical insulation support). In addition, the iSCAPE system is also provided with a ruler that is used to measure the distance from the PM feed point as illustrated in Fig 1. The dimensions of the container are 60×20×4 mm (length × width × height). The power supply can provide a voltage of up to 20 kV. The mineral oil has a viscosity of 14.2-17.2 cSt (11.9-14.5 mPa*s) and density of 0.84 g/mL at 25 °C.

**Fig 1.**
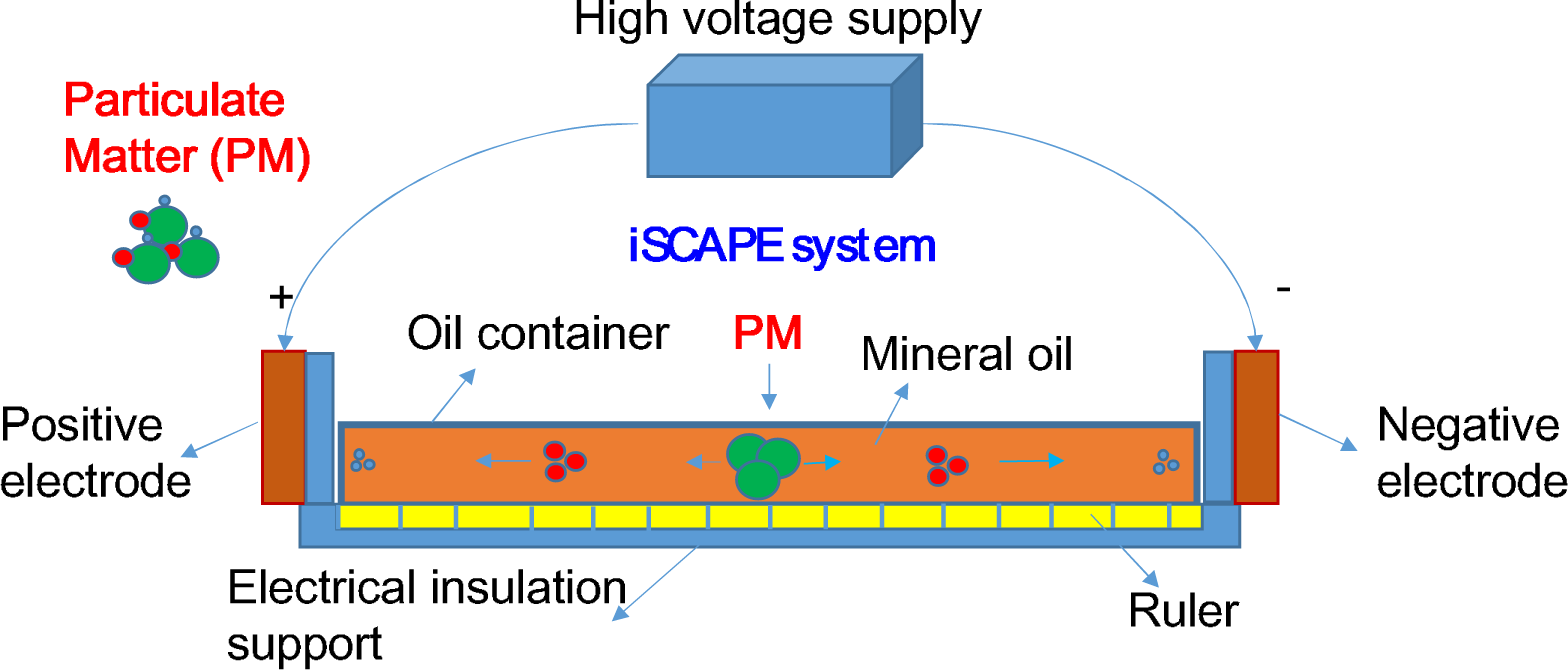
Experimental setup for fine sieving of atmospheric particles using mineral oil electrophoresis (iSCAPE) powered by a high voltage supply together with a mineral oil. The dimensions of the container are 60×20×4 mm(length × width × height).

### iSCAPE of atmospheric particles of various sizes

To test the iSCAPE system, we used atmospheric samples previously collected using automobile air conditioning filters for Beijing, Zurich, and San Francisco *(16)*. Here, we also collected Beijing’s soil and road dust samples for testing the system. When operating the iSCAPE, approximately 1 mL mineral oil was first added into the electrophoresis container. Secondly, the power supply with desired voltage was turned on until being stable without air breakdown between the two electrodes. Lastly, approximately 20 μL mineral oil suspension with the tested samples dissolved (atmospheric particulate matter, soil sample or road dusts) was pipetted into the oil container from the sample feeding point as illustrated in Fig 1. Depending on the experimental objectives, the tests could last from seconds to minutes to sieve particles or extract desired size particles from the sample mixture. Under the used experimental conditions, particles with different electrical mobility would travel a different velocity in the mineral oil, thus ending up in different locations away from the sample feeding point. For particles with positive charges, they moved toward the negative electrode, while particles with negative charges moved toward the opposite.

### Analysis of iSCAPEd samples

In this work, for different samples, we took samples from various points away from the sample feed points, e.g., 0.5, 1, 1.5, 2 and 3 cm. The samples were further subjected to microscopic analysis using a microscope (BX 63, Olympus Co., Tokyo, Japan). In addition, using a slightly modified iSCAPE system, the particle electrophoresis was also directly conducted on a microscopic slide (S2112, Matsunami Co., Osaka, Japan) such that particles at different points between the electrodes can be continuously imaged using the microscope (corresponding videos are provided as Supporting Information; use of the microscopic slide however could impact the original particle charge distribution in the sample). For particles retrieved from different points between two electrodes, various analyses were conducted. Here, as an example analysis we have calculated their electrical mobility, performed microscopic imaging, and bacterial ATP measurements. The particle electrical mobility was calculated using the following equation *(17)*:

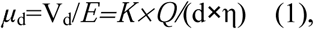

where *μ*_*d*_ is the particle electrical mobility (m^2^/(V*s), V_d_ (m/s) is the particle velocity, *E* (V/m) is the uniform electrostatic field, *K* is a constant, *Q* is the particle charge, *d* is the particle diameter, and η is the medium viscosity. Bacterial ATP measurements, as an example analysis for bacterial separation, were performed using a device (SystemSure Plus, Hygiena, Camarillo, CA). For measuring their ATP levels, 5 μL of mineral oil sample retrieved from different locations was taken and analyzed.

### Statistical analysis

We have tested the iSCAPE system using different samples (atmospheric PM, soil and road dust samples) under different experimental conditions (different electrostatic field strength (3.17 kV and 6.33 kV/cm), different run time (20 s to 6 min). For each sample retrieved, at least five images were taken from different microscopic views. In addition, we have provided videos of imaged particles along the particle moving lines of the mineral oil. Here, mineral oil (microbiology grade) was also imaged to eliminate the possible particle contamination before any experiments as shown in Fig S1.

## Results and Discussion

### Atmospheric particle sieving by iSCAPE under different electrostatic field strength and run time

As shwn in Fig. 2, for particles from different cities iSCAPE has demonstrated different sieving capabilities. For atmospheric samples from Beijing, observed particles seemed to have higher electrical mobility (3.95) compared to those (1.32) of the particles from San Francisco and Zurich. In contrast with the control without the iSCAPE, a large amount of particles travelled to 1.5 cm location from the particle feeding point at a speed of 125 μm/s under the experiemntal conditons tested. As observed in Fig 2, particles with smaller sizes generally moved faster than those larger particles, nonetheless the mobility was proportional to the ratio of particle charge over diameter. Particles from different sources had different particle mobility graphs as shown in the figure under the same iSCAPE operating conditons (3.17 kv/ cm for 2 min). The differences observed for different cities via the iSCAPE were likely due to the different components such as bacteria and metals, and the size distrbutions of their PMs *(5, 16,18)*. For example, air samples from Zurich were shown to have higher fraction of nanoscale particles than those from Beijing *(5)*. For the sampling points listed above, Fig S2 showed the ATP measurements for Beijing’s samples that were iSCAPEd. As seen in the figure, most of bacteria moved to the postive electrode (56%), concentrating within 1.5 cm range from the particle feed point (B-0). For the negative electrode, about 21% was located within 1 cm range from the feed point (B-0). These data suggest that the iSCAPE system can be also used to separate bacterial particles, and a higher fraction of them were shown carrying negative charges. Apparently, a stronger electrostatic field or longer run time are needed to sieve large particles from Zurich and San Francisco.

**Fig. 2.**
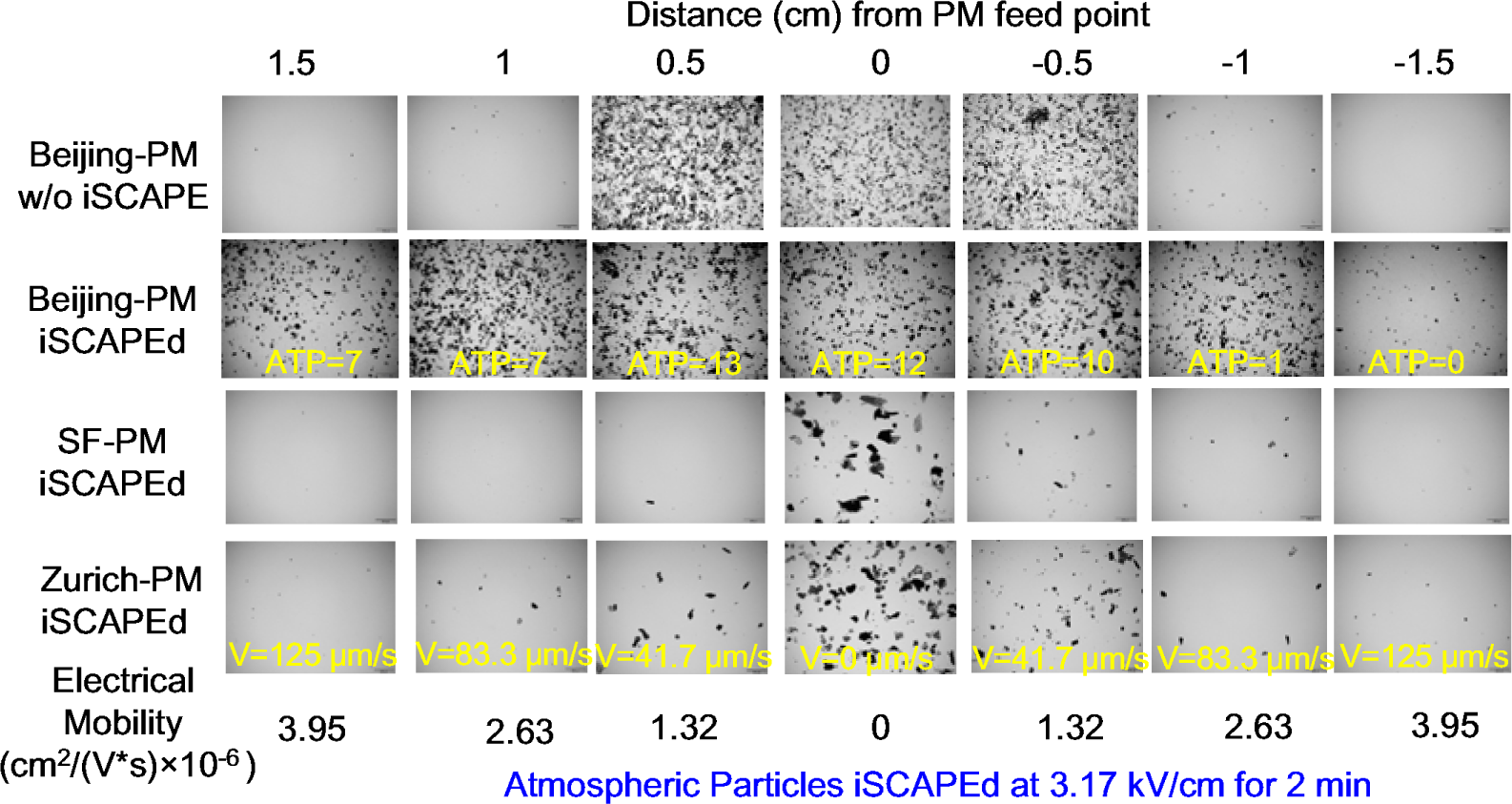
Fine sieving of atmospheric particles collected from three global cities (Beijing, Zurich, and San Francisco(SF)) using the iSCAPE at 3.17 kV/cm for 2 min. Bacterial ATP results for different sampling points are shown in Fig S2. For each sampling point, at least five images (200 μm scale bar) taken from different microscopic views of 10 μL sample.

**Fig. 3.**
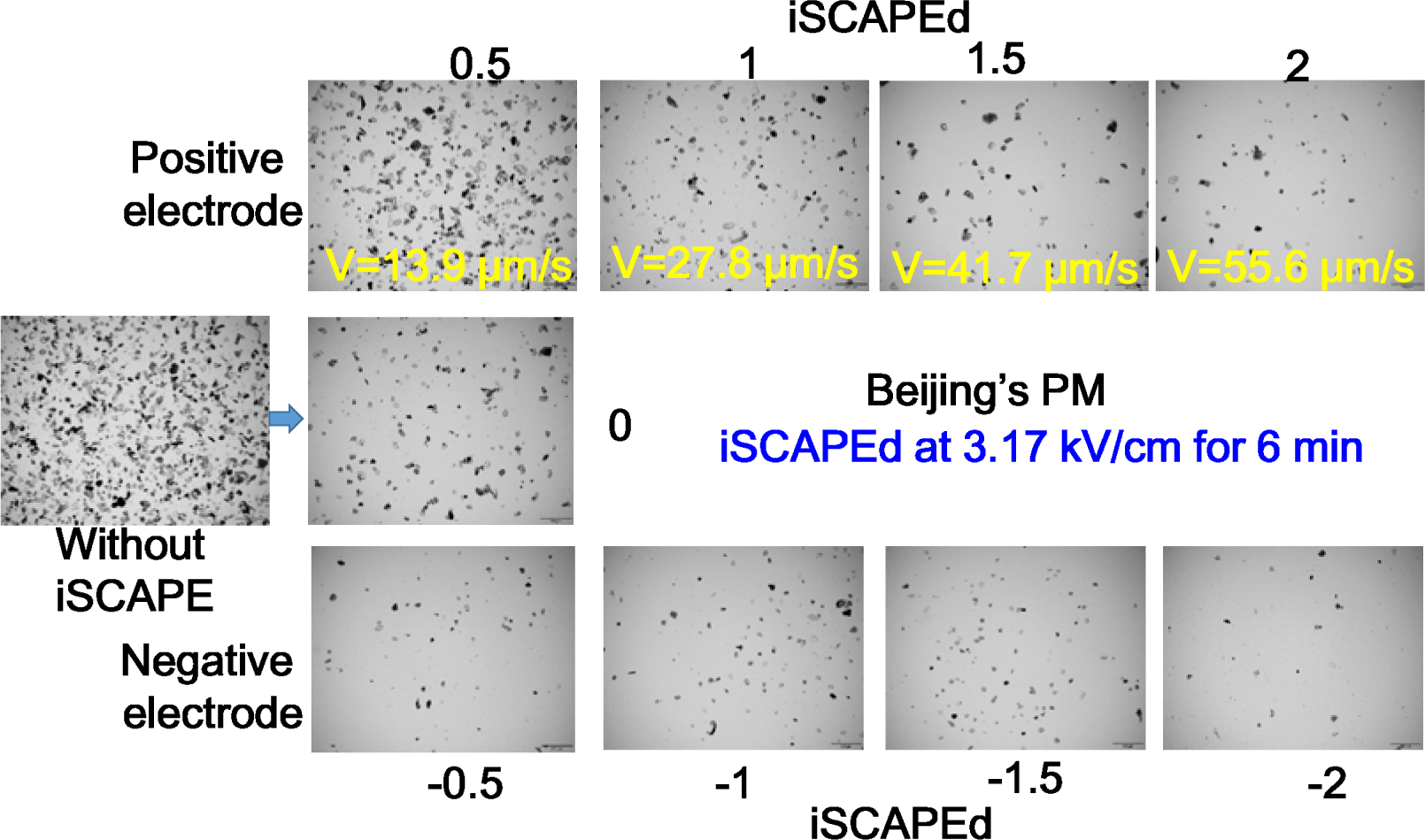
Fine sieving of atmospheric particles from Beijing using the iSCAPE at 3.17 kV/cm for 6 min. Beijing‘s PMs iSCAPEd videos for 3 min with particles along the particle moving line are further provided in File S1 (negative charges) and S2 (positive charges) (Supporting Information). For each sampling point, at least five images (200 μm scale bar) taken from different microscopic views of 10 μL sample.

To further test the iSCAPE capability, we have repeated the test with Beijing’s PM samples but with longer run time, i.e., 6 min, at the same electrostatic field strength (3.17 kV/ cm). Compared to shorter time shown in Fig 2, more particles travelled away from the PM feed point. It can be again seen that particles with smaller sizes generally travelled much faster (55.6 μm/s) than those with larger sizes (13.9 μm/s). In addition to these sampling points, we have provided particle separation information (iSCAPEd for 3 min) along the particle moving line in videos (File S1 and S2, Supporting Information) from which imaged particles can be seen at any locations between the electrodes. Also, as observed from the figure there were more particles carrying negative charges than those carrying negative ones. Data in these videos also demonstrated that the iSCAPE system can efficiently sieve out the particles. Depending on the targets to be obtained, the run time and electrostatic field strength can be fine adjusted.

### Soil and road dust sample sieving by the iSCAPE under different electrostatic field strength and run time

To further validate the iSCAPE system, we have also performed the same tests with Beijing soil and road dust samples (Fig S3, S4, and Fig 4) at 3.17 kV/ cm for 2 min. As observed in Fig S3, and S4 along with videos (File S3 S4, S5, and S6), the iSCAPE system was shown to efficiently sieve soil and road dust particles using their electrical mobility. Again, particles of different sources have demonstrated different mobility graphs under the same operating conditions (the electrostatic field strength and the same run time) given the same viscosity mineral oil. To test high electrostatic field strength, a modified iSCAPE was used, e.g., shorter electrode distance (3 cm) but with the same voltage (19 kV), and the results with Beijing soil sample are shown in Fig 4. As observed from the figure, under higher electrostatic field strength (6.33 kV/ cm), all particles travelled much faster up to 500 μm/s for the location of 1 cm than the lower electrostatic field (3.17 kV/cm), and even within 20 seconds the particles can be well sieved as seen in the figure. There was a clear contrast between samples before and after the iSCAPE test. Images of particles at other particle moving points on the line can be seen in File S7 and S8 (Supporting Information). In addition to air, these data showed that the iSCAPE system can be also applied to many other samples, and the particle sieving can be fine controlled by adjusting the electrostatic field strength and the run time.

**Fig. 4.**
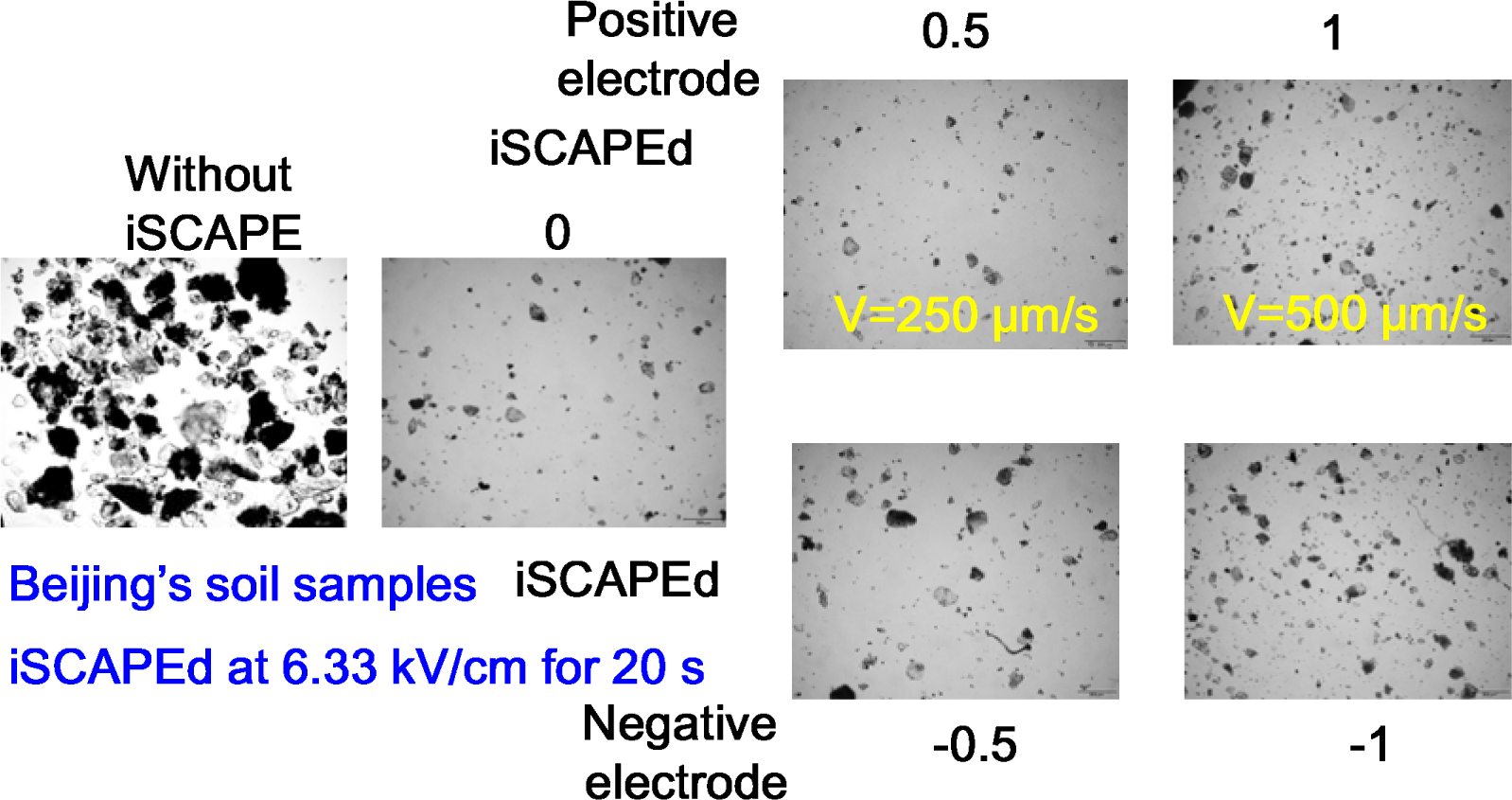
Fine sieving of Beijing’s soil samples using the iSCAPE at 6.33 kV/cm for 20 seconds. Beijing’s soil sample iSCAPEd videos are provided in File S7 (negative charges) and File S8 (positive charges) (Supporting Information). For each sampling point, at least five images (200 μm scale bar) taken from different microscopic views of 10 μL sample.

In this work, we report an invention (the iSCAPE) that can be used to fine sieve, enrich and extract desired particles including bacteria, fungi, pollen and viruses, out of a particle mixture based on their electrical mobility. For particle health or haze formation mechanism study, the iSCAPE can be used to extract selected particles from an air sample using pre-determined operating parameters. The iSCAPE system also holds an immense potential in separation and purification of protein and chemical molecules from a biological sample. In principle, the iSCAPE system can be used to extract any desired targets from a sample of environmental or medical origin, e.g., for an improved PCR detection, by modifying electrical field, mineral oil viscosity, run time and particle electrical mobility. The particle electrical mobility per equation (1) is a function of particle charge, electrophoresis medium viscosity and particle diameter *(17)*. The bacterial particle charge can be attributed to two factors: ionizable groups ((NH_2_) and carboxyl (COOH)) or others present on the cell surface and the external particle frictions *(19)*. To some extent, the latter can be modified by a manual charging process. Therefore, biological and non-biological particles with similar particle sizes could move differently under the same iSCAPE operating condition. The iSCAPE system could be negatively impacted by the moistures in the sample and possible ions in the mineral oil. Certainly, a large amount of work needs to be rapidly explored for the innovative applications of the invented iSCAPE in many different fields such as air pollution, clinical microbiology, and sample purification.

## Acknowledgements

This study was supported by the NSFC Distinguished Young Scholars Fund Awarded to M. Yao (21725701), and the Ministry of Science and Technology (grants 2016YFC0207102). A patent has been submitted for the iSCAPE technology developed here prior to this manuscript submission. M. Yao conceived the research idea, and S. Xu and X. Li performed experiments with equal contributions.

## Supporting Information

**Fig. S1.**
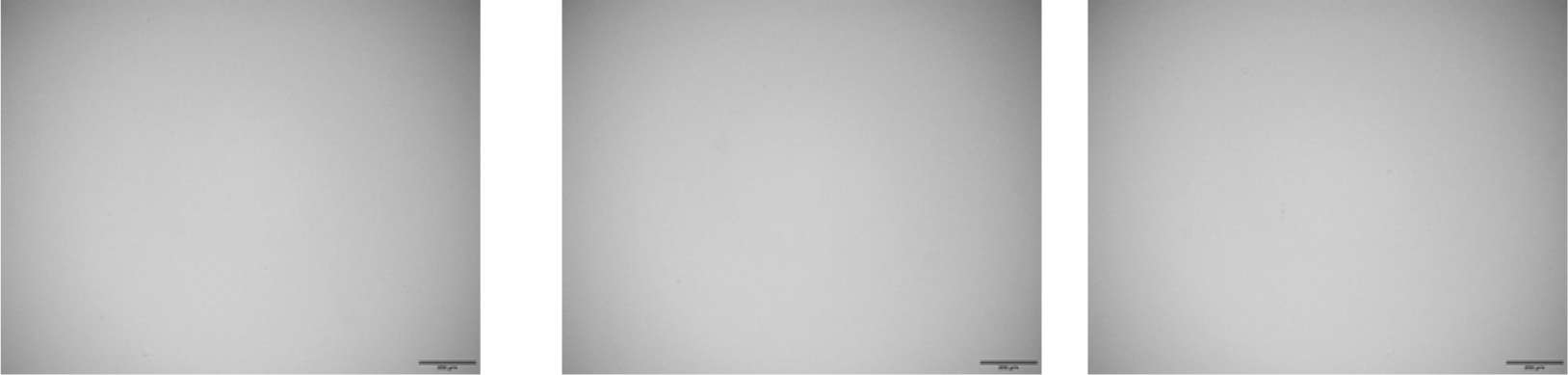
Microscopic images (200 μm scale bar) of mineral oil (microbiology grade) (Sigma-Aldrich) control used for by iSCAPE system: no particle contamination for the mineral oil was detected during the experiments.

**Fig. S2.**
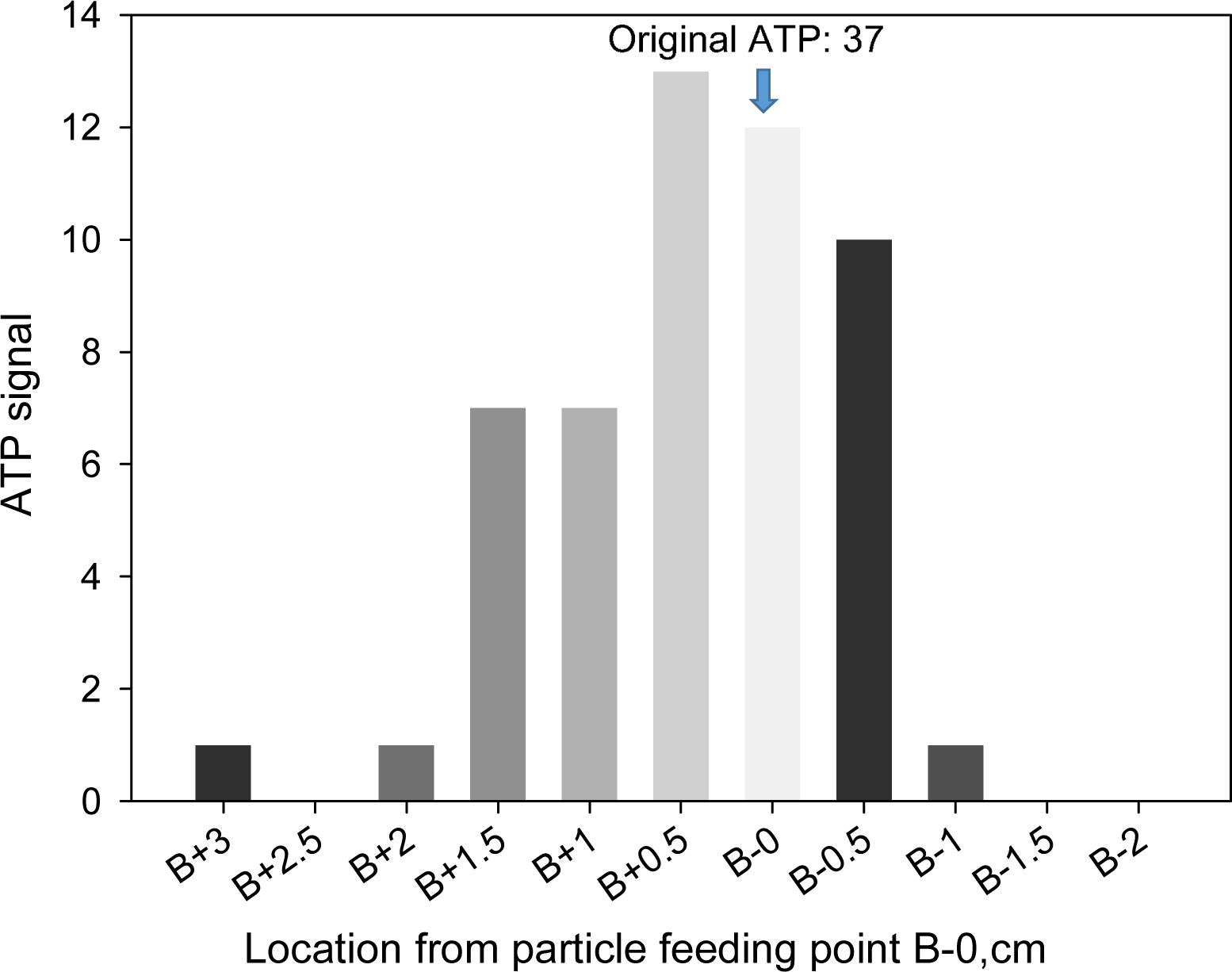
Bacterial ATP detection results for the Beijing’s PM iSCAPEd as shown in Fig 2. Numbers in the figure represent the distances (cm) away from the sample feed point (B-0);and minus and plus signs represent locations toward negative and positive electrodes, respectively. Due to limited sample volume, 5 μL was used for ATP measurement for each sampling point.

**Fig. S3.**
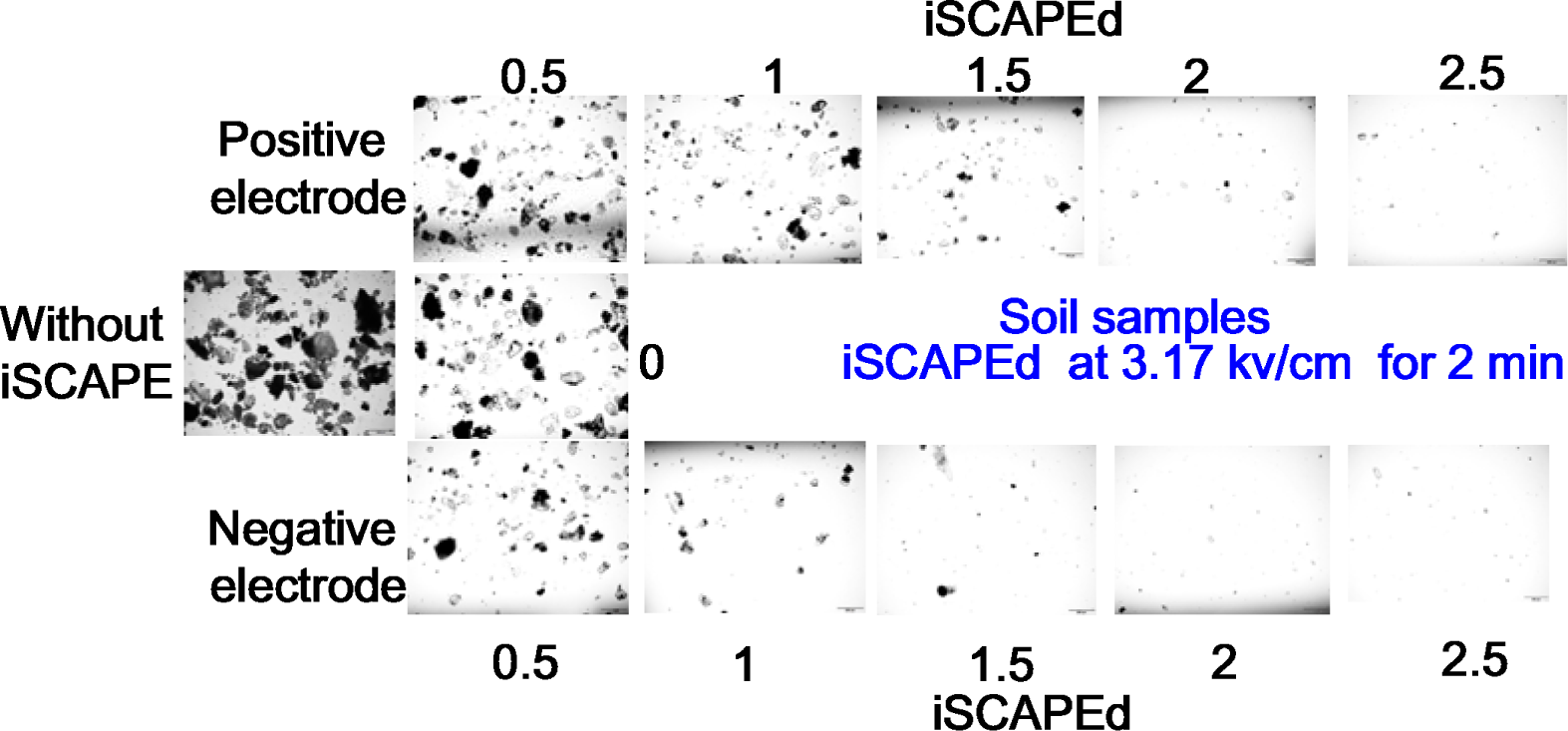
Fine sieving of Beijing’s soil samples using the iSCAPE at 3.17 kV/cm for 2 min. Beijing’s soil sample iSCAPEd videos are provided in File S3 (negative charges) and File S4 (positive charges). For each sampling point, at least five images (200 μm scale bar) taken from different microscopic views of 10 μL sample.

**Fig. S4.**
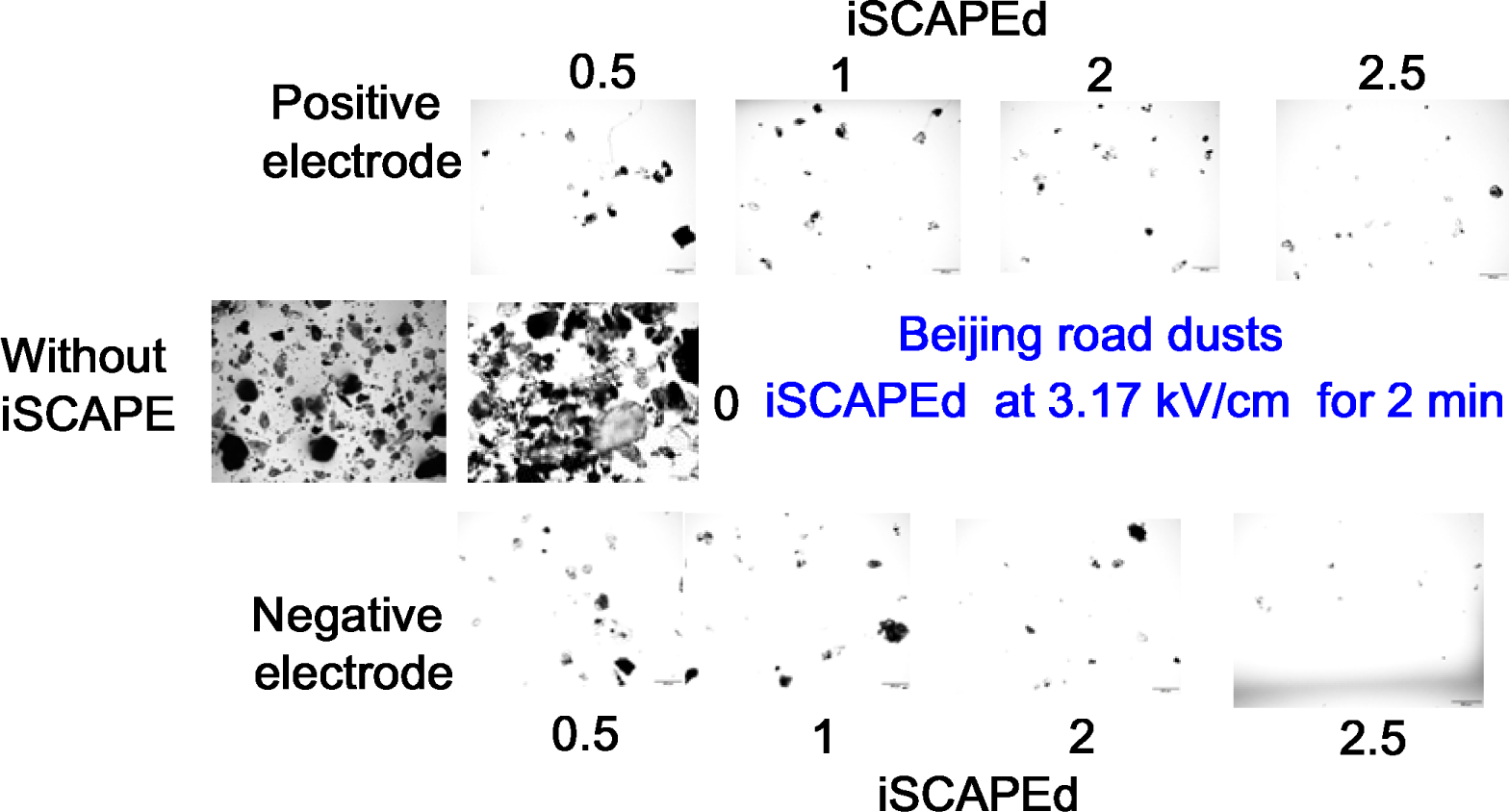
Fine sieving of Beijing’s road dust samples using the iSCAPE at 3.17 kV/cm for 2 min. Beijing’s road dust samples iSCAPEd videos are provided in File S5 (negative charges) and File S6 (positive charges). For each sampling point, at least five images (200 μm scale bar) taken from different microscopic views of 10 μL sample.

## Supporting Files

**File S1** Beijing‘s PMs with negative charges iSCAPEd videos at 3.17 kV/cm for 3 min with imaged particles (200 μm scale bar) along the particle moving line.

**File S2** Beijing‘s PMs with positive charges iSCAPEd videos at 3.17 kV/cm for 3 min with imaged particles (200 μm scale bar) along the particle moving line.

**File S3** Beijing‘s soil samples with negative charges iSCAPEd videos at 3.17 kV/cm for 2 min with imaged particles (200 μm scale bar) along the particle moving line.

**File S4** Beijing‘s soil samples with positive charges iSCAPEd videos at 3.17 kV/cm for 2 min with imaged particles (200 μm scale bar) along the particle moving line.

**File S5** Beijing‘s road dust samples with positive charges iSCAPEd videos at 3.17 kV/cm for 2 min with imaged particles (200 μm scale bar) along the particle moving line.

**File S6** Beijing‘s road dust samples with negative charges iSCAPEd videos at 3.17 kV/cm for 2 min with imaged particles (200 μm scale bar) along the particle moving line.

**File S7** Beijing‘s soil samples with negative charges iSCAPEd videos at 6.33 kV/cm for 20 seconds with imaged particles (200 μm scale bar) along the particle moving line.

**File S8** Beijing‘s soil samples with positive charges iSCAPEd videos at 6.33 kV/cm for 20 seconds with imaged particles (200 μm scale bar) along the particle moving line.

